# Circadian Activity Predicts Breeding Phenology in the Asian Burying Beetle Nicrophorus nepalensis

**DOI:** 10.1101/2024.11.30.626159

**Authors:** Hao Chen, Dustin R. Rubenstein, Guan-Shuo Mai, Chung-Fan Chang, Sheng-Feng Shen

## Abstract

Climate change continues to alter breeding phenology in a range of plant and animal species across the globe. Traditional methods for assessing when organisms reproduce often rely on time-intensive field observations or destructive sampling, creating an urgent need for efficient, non-invasive approaches to assess reproductive timing. Here, we examined three populations of the Asian burying beetle *Nicrophorus nepalensis* from subtropical Okinawa (500 m) and Taiwan mountains (1100-3200 m) that were reared under contrasting photoperiods in order to develop a predictive framework linking circadian activity to breeding phenology. Using automated activity monitors, we quantified adult circadian rhythms and employed machine learning to predict breeding phenology (seasonal versus year-round breeders) from behavior alone. Our model achieved 95% accuracy under long-day conditions using just three behavioural features, and notably, maintained 76% accuracy under short-day conditions when both types are reproductively active, revealing persistent behavioural differences between breeding strategies. These results demonstrate how integrating behavioural monitoring with machine learning can provide both a rapid, scalable method for tracking population responses to climate change and novel insights into species’ adaptive responses to shifting seasonal cues across different elevational gradients in their native range.

## 1. Introduction

In an era of rapid climate change, shifts in breeding phenology have become one of the most conspicuous biological responses to global warming [1]. Such phenological shifts, particularly in reproductive timing, represent a fundamental adaptation that enables species to synchronise reproduction with optimal environmental conditions [2]. This adaptive timing mechanism integrates multiple environmental cues, from photoperiod, temperature [3], and precipitation [4] to snow cover [5]. Yet, climate change is disrupting the reliability of these environmental signals, triggering widespread shifts in reproductive timing across taxa [6], with potentially profound implications for population persistence.

Traditional methods for assessing breeding phenology often present significant challenges. For example, field studies monitoring insect breeding activities along elevation gradients can require extensive resources and prolonged observation periods of up to 19 days per breeding event [7]. Alternative approaches using histological examination of reproductive organs, while precise, are inherently destructive and preclude longitudinal monitoring [8–10]. These methodological constraints have limited our ability to effectively track rapid phenological responses to environmental change.

Behavioural rhythms offer a promising alternative indicator of reproductive state. Across diverse taxa, circadian activity patterns show consistent relationships with breeding conditions. For example, polygynous water skinks (*Eulamprus heatwolei*) display increased movement and social interactions during breeding periods, correlating strongly with reproductive success [11]. Alpine chamois (*Rupicapra rupicapra*) shift from unimodal to multimodal daily activity patterns during breeding seasons [12–14], while Arctic-breeding shorebirds exhibit distinct activity signatures during reproduction [15]. These synchronised behavioural adaptations reflect the optimization of reproductive timing across temporal niches [16, 17].

Recent advances in machine learning have created new opportunities to leverage these behavioural indicators for reproductive monitoring. While machine learning has successfully characterized mammalian behavioural patterns [18] and emotional states [19], its application to reproductive phenology remains largely unexplored. Here, we demonstrate how integrating automated activity monitoring with machine learning can reliably predict reproductive seasonality in populations of the Asian burying beetle (*Nicrophorus nepalensis*). By combining high-resolution behavioural data with controlled breeding experiments, we developed a random forest model that can accurately distinguish seasonal from year-round breeders based only on circadian activity patterns. This non-invasive approach enables unprecedented temporal and spatial resolution in tracking breeding phenology, offering a powerful new tool for monitoring biodiversity responses to environmental change.

## 2. Methods

### (a) Study organism

Burying beetles (Silphidae: *Nicrophorus*) utilize small vertebrate carcasses for both reproduction and provisioning offspring [20]. Post-mating, pairs prepare carcasses by removing hair [21] and applying antimicrobial secretions [22]. The processed carcass is then shaped into a brood ball, buried, and surrounded by eggs laid by the female. After approximately 14 days, third-instar larvae disperse for pupation. Following a 1.5-month pupal period, emerged adults require 3-4 weeks of feeding to achieve sexual maturity [23] before initiating subsequent breeding cycles.

Despite being stenothermic and cold-adapted [24], *N. nepalensis* maintains a broad distribution across Asia [25]. Populations exhibit distinct breeding seasons across their range, with reproductive behavior and ovarian development primarily regulated by photoperiodic cues [26]. This interpopulation variation in breeding phenology reflects local adaptation in reproductive photoperiodism [26].

### (b) Experimental procedures

To investigate adaptive strategies across environmental gradients, we selected three natural populations of *N. nepalensis* from distinct latitudinal and elevation locations: subtropical lowland Okinawa Island, Japan (26.69°N, < 500 m), mid-elevation Mt. Yangming, Taiwan (25.18°N, < 1100 m), and high-elevation Mt. Hehuan, Taiwan (24.14°N, < 3200 m). Previous research revealed that while the high-elevation Mt. Hehuan population remains active year-round [26], lower-elevation populations in Okinawa and Mt. Yangming exhibit distinct winter activity peaks (Shen et al., unpublished data).

We collected beetles using hanging pitfall traps baited with decomposing pork. Wild-caught adults were the used to establish laboratory breeding lines, with experimental subjects comprising both initial generation (wild-type; offspring of wild-caught parents) and subsequent laboratory generations. Third-instar larvae were randomly assigned to either short-(10L:14D) or long-day (14L:10D) photoperiodic treatments throughout pupation (1.5 months pre-emergence) and sexual maturation (1 month post-emergence). Based on previous findings [26] demonstrating that winter-active seasonal breeders maintain their non-reproductive state under long-day conditions, all activity measurements and breeding trials were conducted under long-day conditions (14L:10D).

### (c) Locomotor activity measurement

We quantified circadian activity using the Locomotor Activity Monitor 25 system (LAM25, TriKinetics Inc., Waltham, MA, USA) [27]. Each monitor contained 32 channels equipped with transparent glass tubes (PGT 25 × 125 mm, TriKinetics Inc.) (figure S1a), surrounded by three pairs of infrared emitter-detector gates. Activity was recorded when beetles interrupted these infrared beams. Fresh superworm *Zophobas morio* larvae were provided as food source *ad libitum* at one end of each tube (figure S1b), with the opposite end connected to a 320 ml transparent plastic container filled with soil for refuge (figure S1c). Activity data was transmitted via the Power Supply Interface Unit (PSIU9, TriKinetics Inc.) and collected in 1-minute bins using DAM-System3 software (TriKinetics Inc.). Following a 24-hour acclimation period, activity was monitored continuously for 63 hours under controlled conditions (14L:10D; 16±3°C; RH:83-100%) in a walk-in growth chamber, with the dark period beginning at 19:00 to minimize external disturbances.

### (d) Breeding type assessment

We assessed reproductive success under different photoperiod treatments (10L:14D versus 14L:10D) across all three populations to determine breeding type. Following activity measurements and a minimum 24-hour rest period, beetles were paired for breeding trials under standardized conditions (14L:10D; 16±3°C; RH:83-100%; Hipoint growth chamber). Each trial used unique male-female pairs in transparent breeding boxes (21 cm × 13 cm × 13 cm) containing 10 cm of soil and a 75±7.5 g fresh mouse carcass. After two weeks, breeding success was assessed by the presence of third-instar larvae. Populations successfully breeding under only one photoperiod treatment were classified as seasonal breeders, while those maintaining high reproductive success under both short- and long-day treatments were designated as year-round breeders.

### (e) Data analysis

Due to complete separation in breeding trials (where all pairs either succeeded or failed within certain treatments), we employed a Bayesian Generalized Linear Model (*bayesglm* function, *arm* R package) with Tukey pairwise comparisons to analyse differences in reproductive success across populations and photoperiod treatments. We then trained a Random Forest Classifier [28] using locomotor activity data (section (c) Locomotor activity measurement) as explanatory variables and breeding type classifications (section (d) Breeding type assessment) as response variables. This approach efficiently handles complex time-series features while avoiding distributional assumptions and multicollinearity issues common to traditional models [29]. We supplemented this analysis with SHapley Additive exPlanations (SHAP) to enhance model explainability.

Activity data were separated into short- and long-day datasets to examine the circadian patterns’ predictive power for breeding type. Raw activity data underwent preprocessing using a sliding window approach and logarithmic transformation to smooth signals and reduce noise [30]. Datasets were randomly split into training (70%) and testing (30%) sets. Using the *tsfresh* Python package [31], we initially extracted nine common features (*extract_features function* with *default_fc_paramter = MinimalFCParameters()*), including sum values, mean, median, minimum, maximum, root mean square, variance, standard deviation, and length. Since three features (median, minimum, and length) showed no variance across individuals (median and minimum = 0, length = measurement duration), they were excluded from further analysis.

Using the remaining six features, we trained a preliminary (1^st^) random forest classifier and calculated corresponding SHAP values. To optimize model complexity, we performed hierarchical clustering on these features using the *shap.utils.hclust* function, identifying and eliminating redundant features where clustering tree terminals contained multiple highly correlated variables. We selected representative features from each cluster based on SHAP importance rankings, retaining the most influential feature while removing others providing redundant information. This minimal feature set trained our final (2^nd^) random forest classifier, maintaining consistent parameters and random states with the preliminary model. For comparison, we trained a third classifier using *tsfresh*’s comprehensive feature set (787 features) to evaluate whether our minimal feature set adequately captured behaviourally relevant patterns for discriminating breeding types.

Finally, we compared feature differences between seasonal and year-round breeders under both photoperiodic treatments using Generalized Linear Models (GLM) with Tukey pairwise comparisons. Count-based time series features were analysed using negative binomial regression (*glm.nb* function, *MASS* R package), while continuous features employed linear regression. All analyses were performed using Python v3.8.8 and R v4.3.1.

## 3. Results

Through controlled breeding experiments, we uncovered distinct reproductive strategies among *N. nepalensis* populations from different elevations (Okinawa: 500 m; Mt. Yangming: 1100 m; Mt. Hehuan: 3200 m) that aligned with specific circadian activity patterns. Populations exhibited significant variation in their reproductive responses to photoperiod (Bayesian GLM, Population × Treatment, χ²₁ = 12.36, *p* = 0.002, table 1a). Both Okinawa and Mt. Yangming populations successfully reproduced only when exposed to short-day conditions during development (Okinawa: *p* < 0.001, figure 1a; Mt. Yangming: *p* = 0.001, figure 1b; table 1b), characterizing them as seasonal breeders. In contrast, the Mt. Hehuan population maintained consistently high reproductive success regardless of photoperiod (*p* = 1.00, figure 1c, table 1b), exhibiting a year-round breeding strategy.

**Figure 1.**
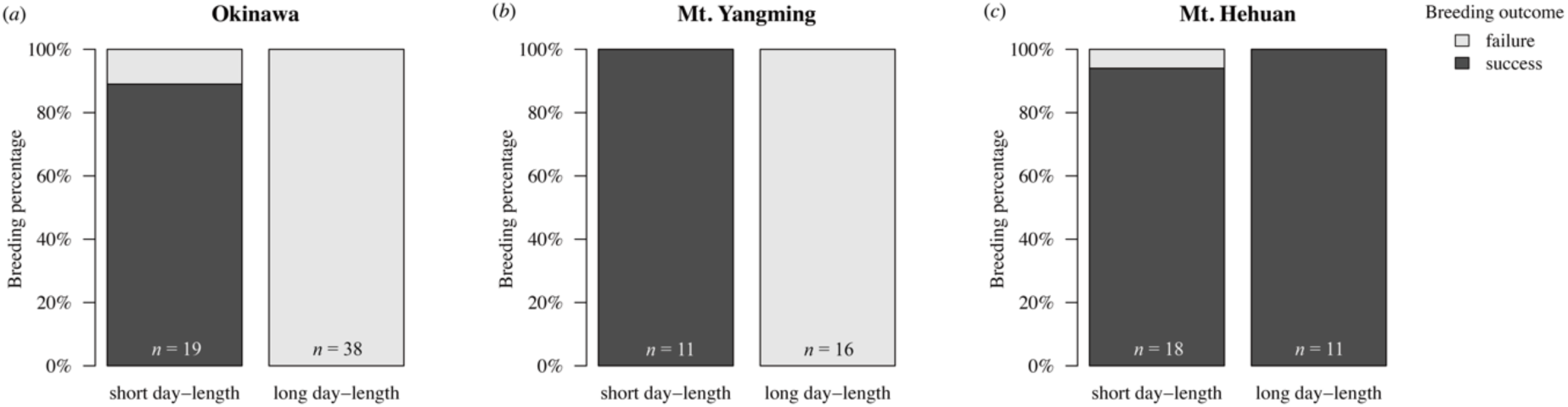
Breeding success of *N. nepalensis* populations in the lab that originated in (a) Okinawa, Japan (500 m), (b) Mt. Yangming, Taiwan (1100 m), and (c) Mt. Hehuan, Taiwan (3200 m) under contrasting photoperiod treatments. Stacked bars represent the proportion of successful (dark grey) and failed (light grey) breeding pairs. Sample sizes (*n*) for breeding pairs are indicated at the base of each bar. Breeding success is defined by the successful emergence of third-instar larvae.

**Table 1.** Population-specific reproductive responses to developmental photoperiod treatments in laboratory-reared *N. nepalensis*. (a) The effects of population origin, photoperiod treatment, and their interaction on breeding success across three populations. (b) Pairwise comparisons of breeding success between individuals developed under short- (10L:14D) or long-day (14L:10D) conditions. Significant effects are in bold.

| Variable | $\chi^2$ | df | P |
| --- | --- | --- | --- |
| Population | 46.19 | 2 | < **0.001** |
| Treatment | 79.31 | 1 | < **0.001** |
| Population × Treatment | 12.36 | 2 | **0.002** |

$n = 113$ (b)
| Contrast | Odds ratio | SE | Z ratio | P |
| --- | --- | --- | --- | --- |
| **Okinawa ($n = 57$):** |  |  |  |  |
| short day-length – long day-length | 1409.11 | 2904.5 | 3.52 | < **0.001** |
| **Mt. YangMing ($n = 27$):** |  |  |  |  |
| short day-length – long day-length | 2656.02 | 6442.3 | 3.25 | **0.001** |
| **Mt. Hehuan ($n = 29$):** |  |  |  |  |
| short day-length – long day-length | 1.00 | 1.3 | -0.003 | 1.00 |

To investigate the relationship between circadian activity patterns and breeding strategies, we monitored locomotor activity of 226 beetles across the three populations (Okinawa: n = 114; Mt. Yangming: n = 54; Mt. Hehuan: n = 58) under long-day conditions (14L:10D) using our automated activity monitoring system (LAM25). Hierarchical clustering analysis of these behavioural data revealed three distinct clusters (figure 2a): one cluster comprising highly correlated measures of mean and total activity counts, another cluster containing root mean square, standard deviation, and variance of per-minute activity counts, and a third cluster containing only maximum counts per minute. From the first two clusters, we selected mean and root mean square as representative features, as they showed the highest importance rankings within their respective clusters in our preliminary random forest model. Given the complex nature of behavioural data, as well as the potential non-linear relationships between activity patterns and breeding strategies, we employed machine learning to determine how each behavioural feature contributes to the model’s predictions and to ensure biological interpretability.

**Figure 2.**
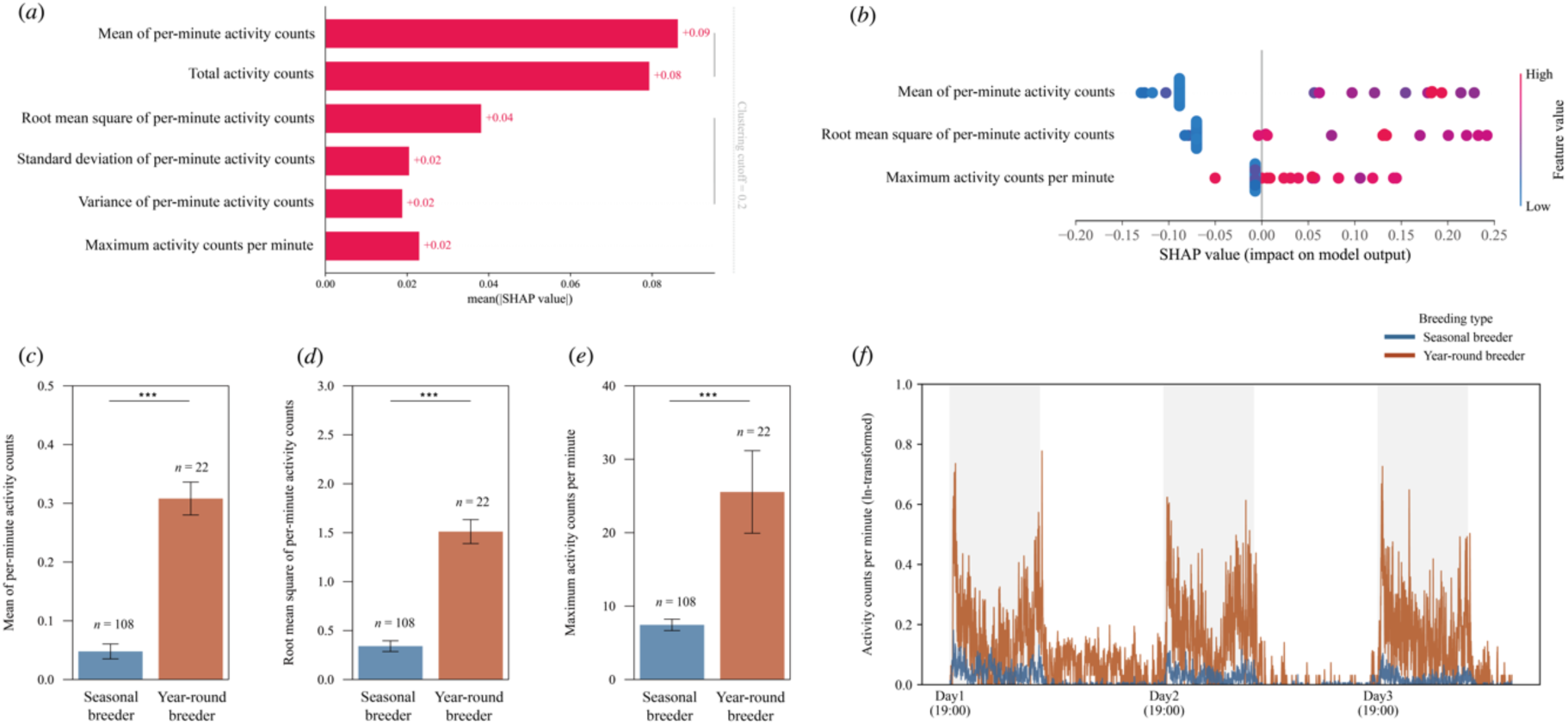
Activity patterns distinguish seasonal from year-round breeders under long-day treatments. (a) SHAP summary plot showing relative importance (mean SHAP value) of activity features in predicting breeding type. Hierarchical clustering tree reveals three distinct feature groups: maximum activity counts per minute provides independent information, while mean of per-minute activity counts and total activity counts form a second correlated cluster, and root mean square, standard deviation, and variance of per-minute activity form a third cluster. Representative features with highest mean SHAP values were selected from each cluster. (b) SHAP beeswarm plot illustrating feature contributions to model predictions, where each point represents an individual beetle (red = high values, blue = low values). SHAP values > 0 indicate stronger prediction for year-round breeders, whereas SHAP values < 0 indicate seasonal breeders. (c-f) Breeding type comparisons showing: (c) mean of per-minute activity; (d) root mean square of per-minute activity; (e) maximum activity counts per minute; and (f) daily activity patterns. Error bars represent standard error. *p* < 0.1, \**p* < 0.05, \*\**p* < 0.01, \*\*\**p* < 0.001.

Using these three representative features—mean, root mean square and maximum counts per minute—we developed a streamlined random forest classifier that achieved exceptional predictive accuracy (accuracy = 0.949, F1 score = 0.949). Remarkably, this minimal feature set achieved comparable performance to a comprehensive model incorporating 787 activity features (accuracy = 0.949, F1 score = 0.949). SHAP analysis revealed that the mean of activity counts per minute contributed most strongly to the model’s predictions, with higher values of these features typically indicating year-round breeders (figures 2b and e).

The predictive power of activity patterns varied with photoperiod. Under short-day treatments, hierarchical clustering identified three distinct feature groups (figure S2a). A random forest model using the representative features from each cluster (maximum counts per minute, total activity counts, and variance) achieved moderate accuracy (accuracy = 0.621, F1 score = 0.588). However, when incorporating the comprehensive feature set, model performance improved substantially (accuracy = 0.759, F1 score = 0.756). This marked improvement in classification accuracy reveals that while seasonal and year-round breeders may appear to exhibit similar activity patterns under short-day conditions, their behavioural responses remain fundamentally distinct when analysed at a finer scale. This finding suggests that despite both types being reproductively active during winter conditions, they maintain distinct behavioural signatures that reflect their differing life-history strategies. SHAP analysis revealed an unexpected pattern: while higher maximum activity still predicted year-round breeders, higher total activity was associated with seasonal breeders (figure S2b), potentially reflecting their sustained daytime activity (figure S2f).

Traditional statistical analyses (GLM) confirmed significant behavioural differences between breeding types across photoperiodic treatments (Treatment × Breeding type, *p* ≤ 0.001, table S1a-5a). Under long-day treatments, year-round breeders exhibited significantly higher mean (*p* < 0.001, figure 2c, table S1b), root mean square (*p* < 0.001, figure 2d, table S2b) and maximum counts per minute (*p* < 0.001, figure 2e, table S3b) than seasonal breeders. However, under short-day treatments, both breeding types showed comparable levels across all the three representative features (total activity: *p* = 0.98, figure S2c, table S4b; maximum counts per minute: *p* = 0.34, figure S2d, table S3b; variance: *p* = 0.08, figure S2e, table S5b).

## 4. Discussion

This study demonstrates how behavioural monitoring can provide a robust non-invasive method for predicting insect breeding phenology (figure 3). By analysing *N. nepalensis* circadian activity patterns with machine learning, we achieved 94.9% accuracy in distinguishing seasonal from year-round breeders. Even more remarkably, our comprehensive behavioural analysis revealed that these populations maintain distinct activity signatures under short-day conditions (75.9% accuracy), despite both types being reproductively active. This methodological breakthrough addresses a critical challenge in studying phenological responses to climate change, as traditional approaches to assessing reproductive status typically require intensive fieldwork [7] or destructive sampling [9, 10]. The strong association that we observed between circadian activity and reproductive state aligns with growing evidence that behavioural rhythms serve as reliable indicators of reproductive status across taxa. Particularly noteworthy is not only the distinct activity level divergence between seasonal and year-round breeders under long-day conditions, but also their persistent behavioural differences under short-day conditions when both are reproductively active. These behavioural signatures likely reflect deeper adaptations to local temporal niches beyond mere reproductive timing, suggesting that populations have evolved integrated life-history strategies that manifest in their activity patterns regardless of current reproductive state. This finding extends our understanding of how animals optimize their temporal activity patterns to maximize reproductive success, a key aspect of life history evolution [32]. Species have evolved adaptive circadian rhythms that enable them to anticipate environmental changes and time specific responses or activities appropriately [17, 33]. For instance, many animals exhibit predictable daily fluctuations in predation risk and have evolved activity patterns that maximize foraging while minimizing mortality risk [34]. Such temporal adaptations are documented across diverse ecosystems, from aquatic fish [35] to desert rodents [36] and cave-dwelling organisms like harvestman [37], demonstrating fine-scale partitioning of temporal niches. These examples illustrate the universal importance of time as an ecological resource in species’ life history evolution.

**Figure 3.**
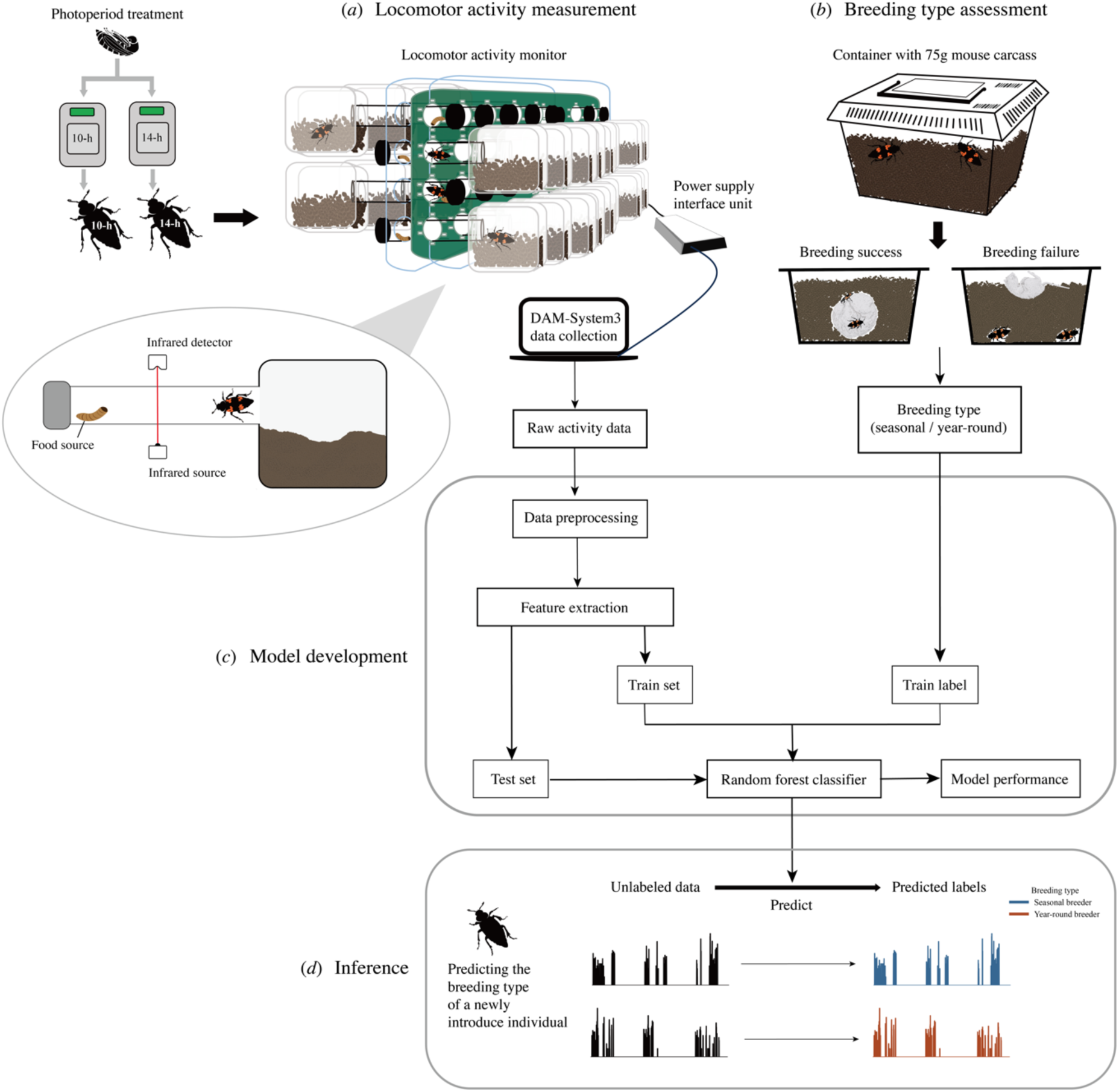
Framework for predicting insect breeding seasonality using circadian activity patterns. Prior to activity monitoring, individuals were randomly assigned to either short- (10L:14D) or long-day (14L:10D) photoperiodic treatments throughout pupation and sexual maturation. (a) Locomotor activity measurement: beetles exposed to different photoperiod treatments are monitored using infrared detectors to record daily activity patterns. Inset shows the experimental setup with food source and shelter area. (b) Breeding type assessment using standardized breeding trials with a 75 g mouse carcass to classify individuals as seasonal or year-round breeders. (c) Model development pipeline showing data processing steps from raw activity data through random forest classifier training. (d) Model application demonstrating how unlabelled activity patterns from newly introduced individuals can be classified into breeding types, enabling rapid assessment of reproductive seasonality. This non-invasive approach allows efficient monitoring of phenological responses to environmental change.

Traditional methods for assessing insect reproductive status have significant limitations. While histological analyses provide accuracy, they are time-consuming and largely restricted to female reproductive organ assessment. As highlighted by Kendra, Montgomery [8] and Tsai, Rubenstein [26] in studies of fruit flies and burying beetles, respectively, determining female ovarian development stages often requires dissection and detailed histological analysis. Even seemingly simpler methods like gonadosomatic index (GSI) depend on sampling timing and often struggle to capture breeding season dynamics [38]. In contrast, our behavioural monitoring approach offers an innovative solution for assessing insect reproductive status. This non-invasive method simultaneously monitors both male and female reproductive behavior, eliminating gender bias inherent in traditional approaches. Furthermore, automated activity recording systems provide unprecedented temporal resolution through continuous data collection, far surpassing the capabilities of traditional field surveys or laboratory dissections. Crucially, behavioural monitoring enables real-time tracking of reproductive state changes, something that is particularly suitable for species with distinct reproductive behaviours like burying beetles. These advantages make our method particularly valuable for studying reproductive responses to photoperiod and temperature changes, as well for assessing climate change impacts on breeding behavior.

As climate change increasingly disrupts seasonal environmental cues [6], understanding and predicting shifts in breeding phenology becomes crucial for conservation. Our machine learning approach, based on easily observable behavioural patterns, provides a scalable tool for monitoring phenological responses across populations and species. This is particularly valuable for tracking rapid evolutionary responses to environmental change, as behavioural adaptations often precede morphological or physiological changes. Looking forward, the integration of automated behavioural monitoring and machine learning opens new possibilities for large-scale phenological studies. This method’s efficiency and non-invasive nature make it especially suitable for long-term monitoring programmes across altitudinal and latitudinal gradients [26], potentially revealing how different populations adjust their reproductive timing in response to environmental pressures. Such insights are crucial for predicting species adaptations to future climate scenarios and developing effective conservation strategies.

## Authors’ contributions

H.C.: data curation, formal analysis, investigation, methodology, visualization, writing—original draft, writing—review and editing; D.R.R.: methodology, writing—review and editing; G.S.M.: formal analysis; C.F.C.: investigation; S.F.S.: conceptualization, data curation, investigation, methodology, project administration, resources, supervision, writing—original draft, writing— review and editing.

## Acknowledgements

We thank Shi-Ping Huang, Shiun-Cheng Chan and Catherine Yung-Yi Lan for their logistical support with the laboratory experiments.

## Supplementary Material

### Supplementary Figures

**Figure S1.**
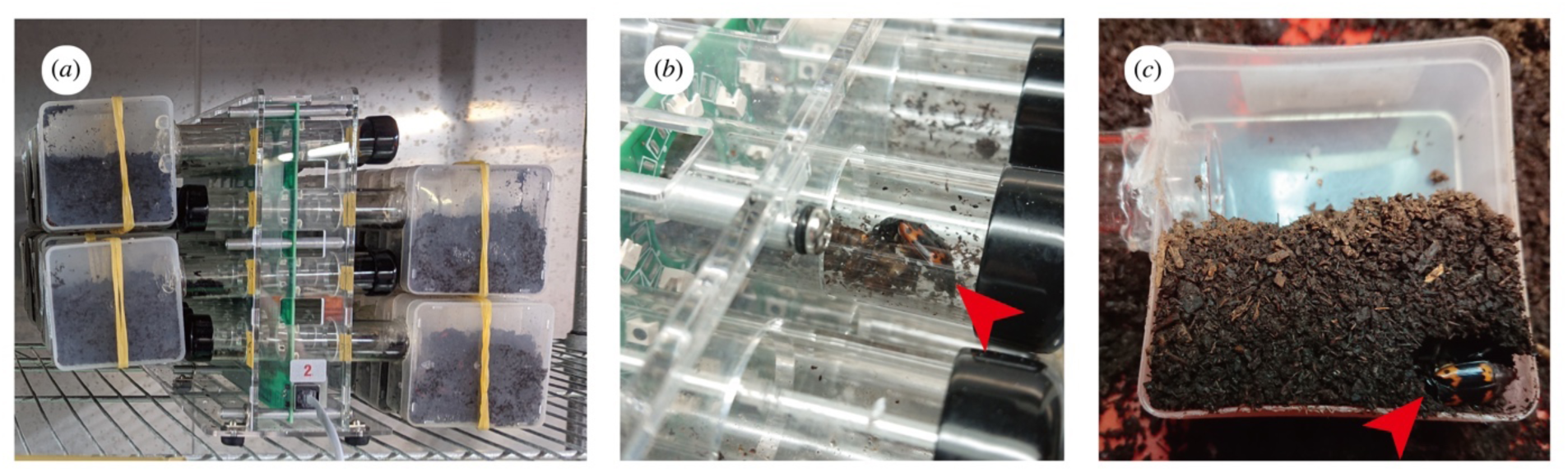
Quantifying circadian activity in *N. nepalensis* using the locomotor activity monitor. (a) Monitor array containing 32 channels with transparent glass tubes for simultaneous individual monitoring. (b) Food source provided *ad libitum* at one end of each tube. (c) Opposite end connected to a 320 ml transparent plastic container filled with soil, serving as refuge. Red arrows indicate beetle positions.

**Figure S2.**
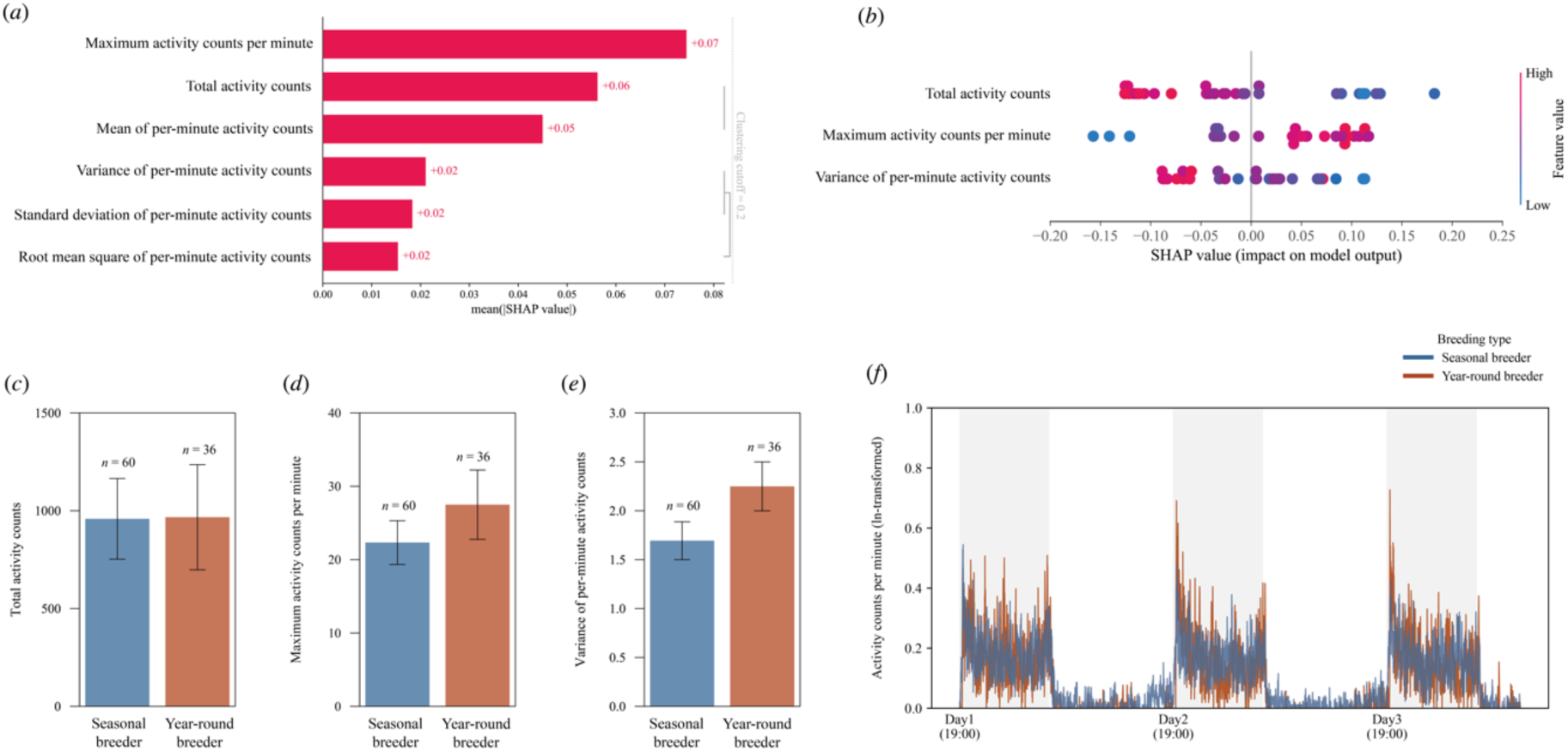
Activity patterns show minimal distinction between breeding types under short-day treatments. (a) SHAP summary plot showing relative importance (mean SHAP value) of activity features in predicting breeding type. Hierarchical clustering tree reveals three distinct feature groups: maximum activity counts per minute provides independent information, while total activity counts and mean per-minute activity counts form a second correlated cluster, and variance, standard deviation, and root mean square of per-minute activity form a third cluster. Representative features with highest mean SHAP values were selected from each cluster. (b) SHAP beeswarm plot illustrating feature contributions to model predictions, where each point represents an individual beetle (red = high values, blue = low values). SHAP values > 0 indicate stronger prediction for year-round breeders, whereas SHAP values < 0 suggest seasonal breeders. (c-f) Breeding type comparisons showing: (c) total activity counts; (d) maximum activity counts per minute; (e) variance of per-minute activity, and (f) daily activity patterns. Error bars represent standard error. *p* < 0.1; \**p* < 0.05; \*\**p* < 0.01; \*\*\**p* < 0.001.

**Table S1.** Effects of photoperiod treatment and breeding types on activity patterns in *N. nepalensis*. (a) The effects of photoperiod treatment, breeding type (seasonal versus year-round breeder), and their interaction on mean of per-minute activity counts. (b) Pairwise comparisons of mean of per-minute activity counts between breeding types under short- (10L:14D) and long-day (14L:10D) treatments. Significant effects are in bold.

| Variable | $\chi^2$ | df | P |
| --- | --- | --- | --- |
| Treatment | 57.96 | 1 | < **0.001** |
| Breeding type | 32.86 | 1 | < **0.001** |
| Treatment × Breeding type | 38.97 | 1 | < **0.001** |

$n = 226$ (b)
| Contrast | Estimate | SE | *t* ratio | P |
| --- | --- | --- | --- | --- |
| **Short day-length ($n = 96$):** |  |  |  |  |
| Seasonal breeder – Year-round breeder | -0.002 | 0.03 | -0.08 | 0.94 |
| **Long day-length ($n = 130$):** |  |  |  |  |
| Seasonal breeder – Year-round breeder | -0.26 | 0.03 | -8.47 | < **0.001** |

**Table S2.** Effects of photoperiod treatment and breeding types on activity patterns in *N. nepalensis*. (a) The effects of photoperiod treatment, breeding type (seasonal versus year-round breeder), and their interaction on root mean square of per-minute activity counts. (b) Pairwise comparisons of root mean square of per-minute activity counts between breeding types under short-(10L:14D) and long-day (14L:10D) treatments. Significant effects are in bold.

| Variable | $\chi^2$ | df | P |
| --- | --- | --- | --- |
| Treatment | 67.94 | 1 | < **0.001** |
| Breeding type | 47.80 | 1 | < **0.001** |
| Treatment × Breeding type | 30.33 | 1 | < **0.001** |

$n = 226$ (b)
| Contrast | Estimate | SE | *t* ratio | P |
| --- | --- | --- | --- | --- |
| **Short day-length ($n = 96$):** |  |  |  |  |
| Seasonal breeder – Year-round breeder | -0.18 | 0.12 | -1.45 | 0.15 |
| **Long day-length ($n = 130$):** |  |  |  |  |
| Seasonal breeder – Year-round breeder | -1.17 | 0.13 | -8.72 | < **0.001** |

**Table S3.** Effects of photoperiod treatment and breeding types on activity patterns in *N. nepalensis*. (a) The effects of photoperiod treatment, breeding type (seasonal versus year-round breeder), and their interaction on maximum activity counts per minute. (b) Pairwise comparisons of maximum activity counts per minute between breeding types under short-(10L:14D) and long-day (14L:10D) treatments. Significant effects are in bold.

| Variable | $\chi^2$ | df | P |
| --- | --- | --- | --- |
| Treatment | 34.35 | 1 | < **0.001** |
| Breeding type | 22.57 | 1 | < **0.001** |
| Treatment × Breeding type | 10.17 | 1 | **0.001** |

*n* = 226 (b)
| Contrast | Odds ratio | SE | Z ratio | P |
| --- | --- | --- | --- | --- |
| **Short day-length (*n* = 96):** |  |  |  |  |
| Seasonal breeder – Year-round breeder | 0.81 | 0.18 | -0.96 | 0.34 |
| **Long day-length (*n* = 130):** |  |  |  |  |
| Seasonal breeder – Year-round breeder | 0.29 | 0.07 | -5.07 | < **0.001** |

**Table S4.**
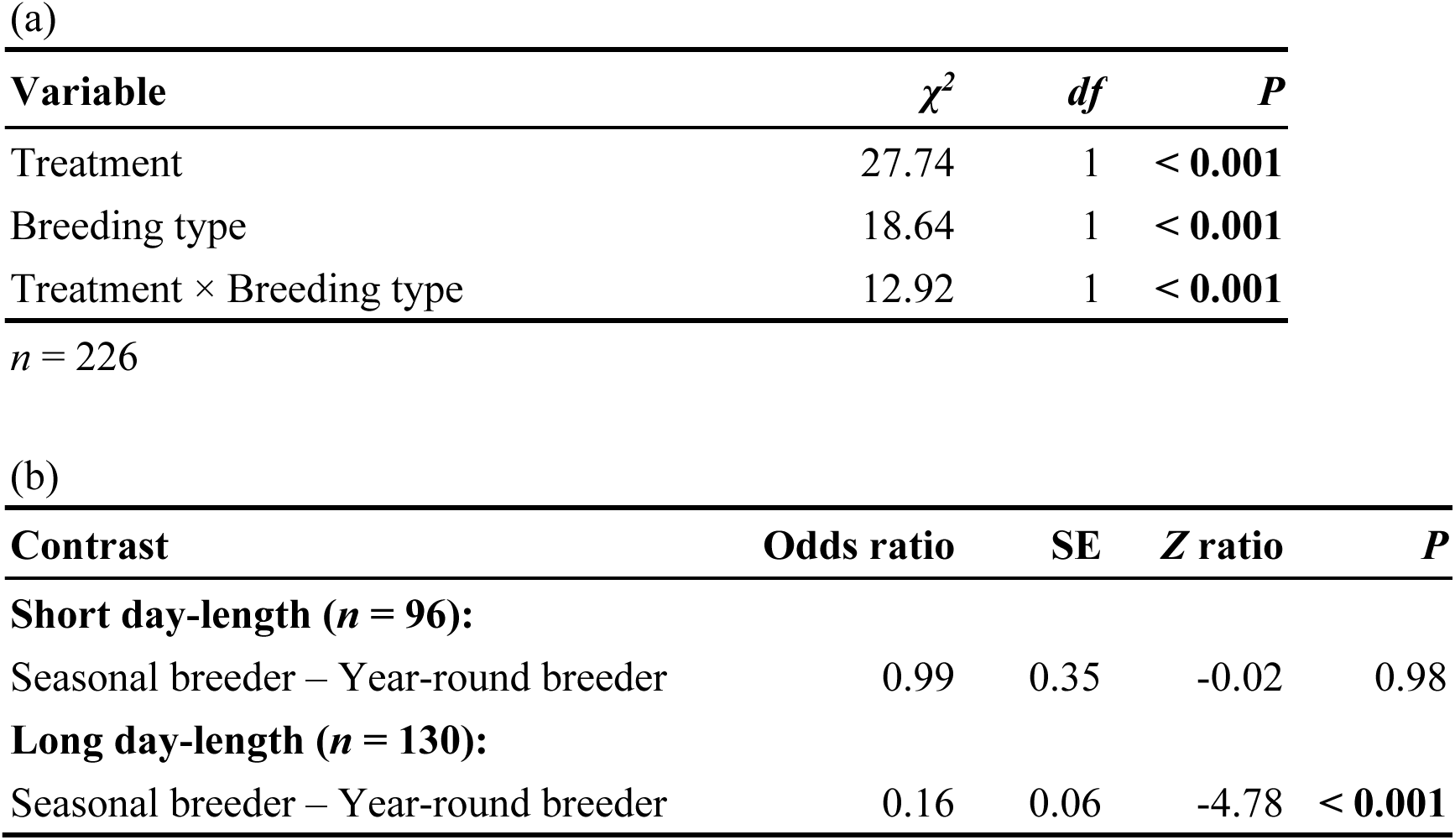
Effects of photoperiod treatment and breeding types on activity patterns in *N. nepalensis*. (a) The effects of photoperiod treatment, breeding type (seasonal versus year-round breeder), and their interaction on total activity counts. (b) Pairwise comparisons of total activity counts between breeding types under short-(10L:14D) and long-day (14L:10D) treatments. Significant effects are in bold.

| Variable | $\chi^2$ | df | P |
| --- | --- | --- | --- |
| Treatment | 27.74 | 1 | < **0.001** |
| Breeding type | 18.64 | 1 | < **0.001** |
| Treatment × Breeding type | 12.92 | 1 | < **0.001** |

$n = 226$ (b)
| Contrast | Odds ratio | SE | Z ratio | P |
| --- | --- | --- | --- | --- |
| **Short day-length ($n = 96$):** |  |  |  |  |
| Seasonal breeder – Year-round breeder | 0.99 | 0.35 | -0.02 | 0.98 |
| **Long day-length ($n = 130$):** |  |  |  |  |
| Seasonal breeder – Year-round breeder | 0.16 | 0.06 | -4.78 | < **0.001** |

**Table S5.** Effects of photoperiod treatment and breeding types on activity patterns in *N. nepalensis*. (a) The effects of photoperiod treatment, breeding type (seasonal versus year-round breeder), and their interaction on variance of per-minute activity counts. (b) Pairwise comparisons of variance of per-minute activity counts between breeding types under short-(10L:14D) and long-day (14L:10D) treatments. Significant effects are in bold.

| Variable | $\chi^2$ | df | P |
| --- | --- | --- | --- |
| Treatment | 14.47 | 1 | < **0.001** |
| Breeding type | 31.55 | 1 | < **0.001** |
| Treatment × Breeding type | 13.05 | 1 | < **0.001** |

*n* = 226 (b)
| Contrast | Estimate | SE | *t* ratio | P |
| --- | --- | --- | --- | --- |
| **Short day-length (*n* = 96):** |  |  |  |  |
| Seasonal breeder – Year-round breeder | -0.56 | 0.32 | -1.75 | 0.08 |
| **Long day-length (*n* = 130):** |  |  |  |  |
| Seasonal breeder – Year-round breeder | -2.26 | 0.35 | -6.44 | < **0.001** |

